# Depletion of BRCA1 Potentiates Progestin-Induced Cytoskeletal Changes in an Ovarian Cancer Cell Model

**DOI:** 10.64898/2026.01.02.697409

**Authors:** Jocelyn Baquier, Noelle E. Gillis, Caroline H. Diep, Laura J. Mauro, Carol A. Lange

## Abstract

Individuals with Hereditary Breast and Ovarian Cancer (HBOC) syndrome carry *BRCA1/2* mutations that predispose them to various cancers. Notably, this condition increases the risk of developing an aggressive ovarian cancer (OC) subtype known as high-grade serous ovarian cancer (HGSOC). The impact of progesterone (P4) on the development of OC subtypes remains unclear. Research suggests a potential interplay between Progesterone Receptor (PR) signaling and BRCA1/2 actions. BRCA1 regulates PR expression and transcriptional activity, while P4 can drive estrogen receptor/PR complexes to BRCA1 DNA binding motifs. Normal fallopian tube tissues from healthy *BRCA1/2* carriers (the cells of origin for HGSOC) exhibit gene signatures similar to those in HGSOC only during the luteal phase of the menstrual cycle, when circulating P4 levels are elevated. This implies a potential role for PR/BRCA complexes in modulating OC initiation and progression. To explore this interaction, BRCA1 was depleted in an ES-2 PR-B+ OC cell models using CRISPR-mediated gene editing. We hypothesized that BRCA1 depletion would alter PR-mediated signaling and reveal novel regulatory mechanisms. Depletion of BRCA1 led to increased total basal PR protein levels and site-specific hyperphosphorylation at S294 and S81 following R5020 (synthetic progestin) treatment. RNA sequencing of NE-1 and KO-BRCA1 pools treated with R5020 identified a high-confidence set of BRCA1-dependent, R5020-induced genes predominately involved with migration and cytoskeletal organization. Transwell assays revealed that R5020 treatment stimulates migration, an effect amplified upon BRCA1 depletion. Inhibition of PIP5K1C using UNC3230 blocked R5020-induced migration in one of the KO-BRCA1 cell pools with pan-Rho inhibitor CT04 showing similar results. The Rac1 inhibitor EHT-1864 suppressed all R5020-induced migration. In contrast, co-treatment with a pan-ROCK1/2 inhibitor Y-27632 resulted in an increase in R5020-induced migration, which was further enhanced with BRCA1 depletion. Overall, these findings establish a crosstalk between BRCA1 and PR signaling, where BRCA1 depletion sensitizes OC cells to progestin-induced migration by altering PR phosphorylation and remodeling cytoskeletal organization via PIP5K1C, Rho-GTPase, and Rac1-dependent mechanisms.

## INTRODUCTION

Hereditary Breast and Ovarian Cancer (HBOC) syndrome is an autosomal dominant inherited disorder where women who carry germline mutations in the *BRCA1* or *BRCA2* genes have a dramatically higher risk of breast and ovarian cancers (1,2). A deleterious germline mutation in these genes results in a cumulative lifetime risk of ovarian cancer (OC) ranging from 11-60% (3,4). Specifically, the risk of malignancy for OC in an individual with a germline *BRCA1* pathogenic mutation is 39-44%, while those with a *BRCA2* pathogenic mutation is 11-17% (5). Clinically, HBOC-associated OC is distinguished by an earlier premenopausal age of onset and a highly aggressive phenotype compared to sporadic cancers (6). High-grade serous ovarian carcinoma (HGSC) is the most lethal ovarian cancer subtype and the predominant form observed in HBOC (7). While tumors in *BRCA1/2* (mBRCA) mutation carriers exhibit platinum sensitivity and high initial response rates to chemotherapy (8), current risk management approaches remain limited in their ability to provide effective prevention or early detection (9).

The BRCA1 and BRCA2 proteins are tumor suppressors that function as primary regulators of the repair of double-stranded DNA breaks via error-free homologous recombination (HR) pathways (10–12). Loss of functional BRCA proteins impairs HR, forcing cells to utilize other error-prone repair pathways, such as non-homologous end joining (NHEJ), which results in cumulative DNA damage, chromosomal instability and ultimately cancer (13,14). Beyond their role in DNA repair, BRCA proteins are also involved in transcriptional control (15), cell cycle progression (16) and cell division (17).

The tissue-specific tumor preference observed in *BRCA1* and *BRCA2* mutation carriers (localized primarily to the breast, ovary, and fallopian tube) strongly implicates a role for steroid receptors (SRs) in HBOC. The functions of SRs and BRCA are intimately connected to regulation of cell cycle mediators and cell proliferation. SR target genes include growth factor receptors/ligands (18,19), cell cycle proteins (20,21), adhesion molecules (22,23) and interestingly, DNA damage repair proteins (24). In breast cancer models, BRCA proteins regulate the expression of steroidogenic enzymes and SRs and also directly inhibit progesterone receptor (PR) transcriptional activity (15,25–27). In addition, normal fallopian tube tissues from healthy *BRCA1/2* carriers, the cells of origin for HGSOC, exhibit gene signatures similar to HGSOC only during the luteal phase of the menstrual cycle, when circulating progesterone levels are elevated (28). This suggests that when BRCA1 is depleted it may lead to progesterone sensitivity.

Intriguingly, recent studies (29) on ER/PR crosstalk have shown that in the presence of estrogen, progesterone can drive estrogen receptor/PR complexes to BRCA1 DNA binding motifs. While existing studies have highlighted the potential interplay between PR signaling and BRCA1/2 actions in breast cancer models, a deeper understanding of how PR and BRCA1 interact in ovarian tissues is crucial for uncovering the specific mechanisms driving the development and metastatic spread of HGSOC.

Our studies described here reveal that BRCA1 serves as a critical regulator of PR signaling in an ovarian cancer cell model. BRCA1 depletion sensitizes OC cells to progestin-induced migration by altering PR phosphorylation and remodeling cytoskeletal organization.

## MATERIALS AND METHODS

### General Reagents

Where applicable, cells were treated with the following reagents at the indicated concentrations: 10nM R5020 (Perkin Elmer, NLP004005MG), 10µM UNC3230 (Med Chem Express, HY-110150), 1µg/ml Rho Inhibitor I (Cytoskeleton, Inc, CT04), 10µM Y27632 dihydrochloride (Med Chem Express, HY-10583), and 25µM EHT1864 (Med Chem Express, HY-16659).

### Cell Lines and Culturing

All cell lines were maintained at 37°C under 5% CO_2_ in cell culture incubators (Eppendorf, 6734010015). ES2 cells were cultured in McCoy’s 5A medium (ATCC, 30-2007) supplemented with 10% charcoal stripped fetal bovine serum (i.e., DCC; Corning, 35072CV), 1% penicillin-streptomycin (i.e., P/S; Gibco, 15070063), and 0.5mg/ml G418 Sulfate (Corning, 61234RG). For experiments involving R5020 treatment, cells were hormone-starved in phenol red-free MEM (Gibco, A1048801), supplemented with 5% DCC, and 1% P/S. All cell lines were routinely tested for mycoplasma using e-Myco PLUS Mycoplasma PCR Detection Kit (Bulldog-Bio, 2523448) and confirmed negative prior to use in experiments.

### Cell Line Generation

The stable cell line, ES2 GFP-PR-B+, was established to express a PR-B isoform tagged at the N-terminus with Green Fluorescent Protein (GFP) as previously described (30). BRCA1 knockout in these parental lines was conducted using CRISPR/Cas9-mediated genome editing, performed by the University of Minnesota Genome Engineering Shared Resource (GESR) at the Masonic Cancer Center. Initial sequencing analyses prior to editing revealed that these parental cells are triallelic for BRCA1 due to triploidy 17 so subsequent screening evaluated the status of all alleles. Single guide RNAs (sgRNAs) were designed to target exon 10 of the *BRCA1* gene to induce loss-of-function mutations. Targeting sequences were XXXXXX and XXXXX. Plasmid constructs were transfected into cells using FuGene and, following editing, cells were subjected to selection and isolated as single-cell–derived clones. Individual clones were expanded and screened for successful *BRCA1* knockout using molecular analyses of the targeted genomic region to detect indels that should disrupt the gene with sequence analysis determine using TIDE software (31). Clones with confirmed triallelic disruption of *BRCA1* were expanded and maintained under standard culture conditions. The extent of BRCA1 expression was confirmed at the protein level along with the associated PR-B protein for each clone, to insure consistent PR-B expression between clones. Based on this protein analysis, single clones were pooled as follows: **Pool A** (KO-A) = clones #15A, 44, 80. **Pool B** (KO-B) = #1, 15B. Non-edited clones include: NE-1 = parental line; NE-2 = clone WT18. Following creation and expansion of pools, expression of BRCA1 and PR-B was re-confirmed for the NE clones and KO-BRCA1 pools, and lines frozen and stored for future experiments.

### Immunoblotting

Nuclear extractions were obtained using the NE-PER Nuclear and Cytoplasmic Extraction Kit (Thermo Scientific, 78835), quantified using the Bio-Rad Protein Assay (Bio-Rad, 5000006), and equal protein concentrations were resolved on 7.5% SDS-PAGE gels. Alternatively, proteins from whole cell lysates were harvested using radioimmunoprecipitation (RIPA) lite lysis buffer [0.15 M NaCl, 6 mM Na_2_HPO_4_, 4 mM NaH_2_PO_4_, 2 mM EDTA, 0.1 M NaF and 1% Triton-X 100 in H_2_O supplemented with 1X complete mini protease inhibitors (Roche, 11836153001), 1X PhosSTOP tablet (Roche, 4906837001), 25 mM β-Glycerophosphate (BGP), 1 mM Phenylmethanesulfonylfluoride (PMSF), 20 µg/ml aprotinin (Fisher Bioreagents, BP250310), 5mM NaF and 0.05 mM Na_3_VO_4_]. Lysates were cleared via centrifugation, quantified using the Bio-Rad Protein Assay, and equal protein concentrations were resolved on 8% SDS-PAGE gels.

Proteins were transferred to Immobilon-P polyvinylidene difluoride membranes (Millipore Sigma, IPVH00010). Membranes were blocked for 1 hour at room temperature in phosphate-buffered saline/0.1% Tween-20 (PBST) containing 5% dried milk. Western blots were probed overnight at 4°C in PBST containing 1% milk using the following primary antibodies: BRCA1 (D-9) (Santa Cruz, sc-6954), PR (F-4) (Santa Cruz, sc-166169), phosphor-Ser294 PR (Custom: (32)), phosphor-Ser345 PR (Custom: (20)), phosphor-Ser190 PR (Custom: (33)), phospho-Ser81 PR (Custom: (34)). HDAC2 (C-8) (Santa Cruz, sc-9959) and GAPDH (Santa Cruz, sc-47724) were used as loading controls for nuclear fractions and whole cell lysates, respectively. As appropriate, HRP-conjugated goat anti-rabbit (Bio-Rad, 170-6515) and goat anti-mouse (Bio-Rad, 170-6516) secondary antibodies were used to detect their respective primary antibodies.

Membranes were developed using WesternBright Quantum HRP Substrate (Advansta, K-12042) or SuperSignal West Pico Plus Chemiluminescent Substrate (Thermo Scientific, 34580).

### Real Time Quantitative PCR (RT-qPCR)

Cells were plated in triplicate in 6-well plates. After 24 hours in phenol red-free, charcoal-stripped media, cells were treated with 10nM R5020 or equivalent volume vehicle (ethanol) for 6 or 24 hours. Cells were then washed with PBS and total RNA was extracted using TriPure Isolation Reagent (Roche, 11667165001) and isopropanol precipitation. Using the qScript cDNA SuperMix (Quanta Bio, 101414108), RNA (1.0 µg) was reverse transcribed to cDNA according to the manufacturer’s instructions. qPCR was performed using LightCycler FastStart Essential DNA Green Master SYBR (Roche, 06402712001) on a LightCycler® 96 Instrument (Roche). Human primer sequences were the following: BRCA1 F (5’-ACCTTGGAACTGTGAGAACTCT-3’), R (5’-TCTTGATCTCCCACACTGCAATA-3’); FOXO1 F (5’-ACGAGTGGATGGTCAAGAGC-3’), R (5’-GCACACGAATGAACTTGCTG-3’); PR F (5’-GTCAGTGGGCAGATGCTGTA-3’), R (5’-TGCCACATGGTAAGGCATAA-3’); p21 F (5’-GACTCTCAGGGTCGAAAACG-3’), R (5’-GGATTAGGGCTTCCTCTTGG-3’); TBP F (5’-ACAACAGCCTGCCACCTTAC-3’), R (5’-TGTGGGGTCAGTCCAGTGCC-3’). qPCR cycling conditions were the following: initial denaturation at 95°C for 10 min, denature at 95°C for 10sec, anneal at 60°C for 10 sec, and extension at 72°C for 5 sec for 45 cycles. All gene expression levels were normalized to housekeeping gene TBP.

### Gene Expression Profiling

#### Sample collection and Sequencing

Cells were plated in triplicate in 6-well plates. After 24 hours in phenol red-free, charcoal-stripped media, cells were treated with 10nM R5020 or equivalent volume vehicle (ethanol) for 6 hours. RNA (1.0 µg) was extracted and purified using RNeasy Mini Kit (Qiagen, 74104) according to manufacturer’s instructions. Purity of the total RNA samples was assessed via BioAnalyzer (Agilent) and samples with an RNA integrity score > 8 were used for library construction by the University of Minnesota Genomics Center. Libraries were built from individual samples with the TruSeq Stranded mRNA kit (Illumina). RNA-Seq libraries were pooled and sequenced on the Illumina NovaSeq X Plus with 50 bp paired-end reads. An average of 20 million reads were collected per sample.

#### Data Analysis

Quality scores across sequenced reads were assessed using FASTQC. Illumina adapters were removed using Trim-Galore. For alignment and transcript assembly, the sequencing reads were mapped to hg38 using STAR (35). Sorted reads were counted using HTseq (36) and differential expression analysis was performed using DESeq2 (37). Genes with a p-value of <0.05 and a log2 fold change greater than 1 or less than - 1 were considered differentially expressed. Comparative gene set enrichment analysis (GSEA) was performed using the ClusterProfiler package (38) to identify differentially enriched biological functions between experimental conditions. Ingenuity Pathway Analysis (IPA, Qiagen) was used to determine enriched canonical pathways and upstream regulators between experimental conditions.

#### Migration Assays

Cells were plated in 10cm plates. After 24 hours in phenol red-free, charcoal-stripped media, cells were treated with 10nM R5020 or equivalent volume vehicle (ethanol) for 24 hours. Cells were then harvested using StemPro™ Accutase™ Cell Dissociation Reagent (Gibco, A1110501), washed and resuspended in serum-free media (SFM) composed of phenol red-free MEM supplemented with 1% P/S. Cells (50k) were seeded in triplicate in 8.0 µm pore 24-well transwell inserts (Falcon, 353097) along with the following treatments: 10nM R5020, 10nM R5020 + 10µM UNC3230, 10nM R5020 + 1µg/ml CT04, 10nM R5020 + 10µM Y-27632, 10nM R5020 + 25µM EHT1864, or equivalent volume vehicle (ethanol or dimethyl sulfoxide). 10% FBS or serum-free control medium was placed in the bottom chamber of the 24-well plates along with the respective treatment and the cells were incubated for 6 hours. Membranes were fixed with 4% paraformaldehyde, stained with Harris Hematoxylin Solution (Millipore Sigma, HHS16), and imaged at 10x magnification using a Nikon Ts2 inverted microscope equipped with a DS-Fi2 color camera. The number of migrated cells were counted for five representative fields per chamber and averaged between conditions.

#### Statistical Analysis

Datasets were tested for a normal distribution using the Shapiro-Wilk normality test followed by, as appropriate, a one-way or two-way ANOVA along with the Tukey multiple comparison test (Prism 10; GraphPad Software). Significance was determined at 95% confidence level and data represented show the mean ± standard deviation (n=sample size).

## RESULTS

### KO of BRCA1 in ES2 PR+ cells leads to activation of PR signaling

To validate BRCA1 depletion in ES2 PR-B + CRISPR-KO cell lines, protein expression was quantified via western blot analysis (Figure 1A). Both KO-BRCA1 pools, KO-A and KO-B, exhibited significantly reduced levels of BRCA1 protein expression compared to NE-1 control (p = 0.0030 and p < 0.0001, respectively). While KO-B also showed a significant reduction relative to NE-2 (p = 0.0015), NE-2 demonstrated lower basal total PR protein expression when compared to NE-1. Given the robust PR levels and consistent BRCA1 expression in NE-1, it was selected as the primary control for all subsequent experiments. Furthermore, to validate knockdown at the transcriptional level, *BRCA1* mRNA expression was quantified across all cell lines using RT-qPCR analysis (Figure 1B). There was no significant difference in transcript levels between the NE-1 and NE-2 controls. Both KO-BRCA1 pools, KO-A and KO-B, exhibited a significant reduction in *BRCA1* mRNA expression levels (p < 0.001) when compared to both control lines. These results confirm successful CRISPR-mediated BRCA1 depletion across both pools.

**Figure 1.**
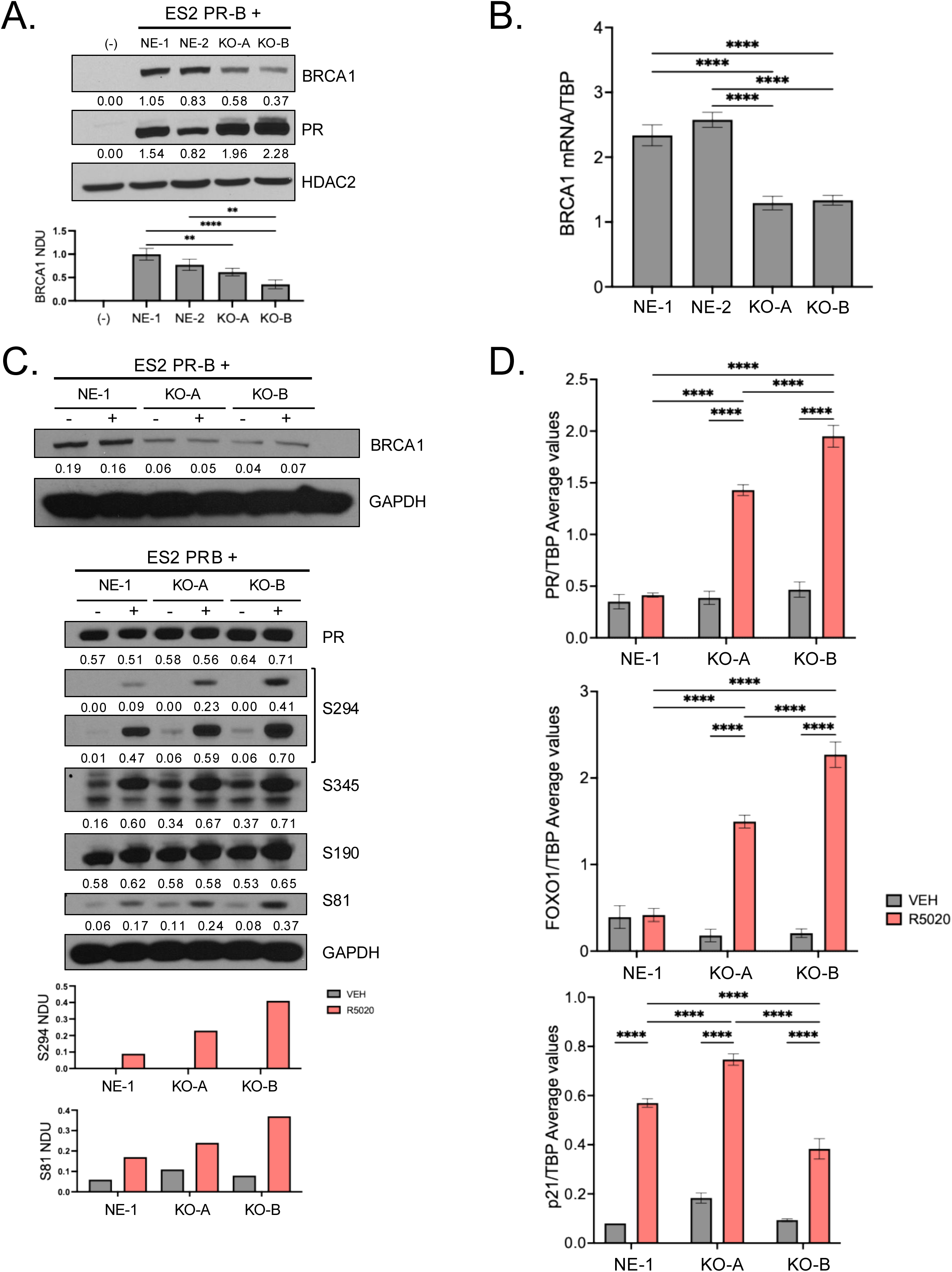
KO of BRCA1 in ES2 PR+ cells leads to activation of PR signaling. **(A)** Basal protein expression of ES2 PR-B+ KO-BRCA1 cell models. Top: Western blot for BRCA1, PR. HDAC2 = loading control. Cell line HCC1937 = negative control for BRCA1. Values under blots represent normalized densitometry units (NDU) for the protein of interest/respective loading control. Graph: NDU for BRCA1 (n=3) shown as mean ± SD, with significance shown as ** p ≤ 0.0030, **** p < 0.0001. **(B)** Basal transcriptional regulation of BRCA1 mRNA. TBP (TATA-box binding protein) = housekeeping gene. Graph: mean ± SD, **** p< 0.0001. **(C)** Top Blot: Western blot of BRCA1. Bottom Blot: Western blot for PR, and phosphorylated PR at serine sites (Ser294, Ser345, Ser190, & Ser81) following treatment with vehicle (ethanol) or 10nM R5020 for 1 hour. GAPDH = loading control. Values under blots represent NDU for the protein of interest/respective loading control. Graph: NDU for S294 (top) and S81 (bottom). **(D)** Transcriptional regulation of PR, FOXO1, and p21 mRNA following treatment with vehicle (ethanol) or 10nM R5020 for 6 hours. TBP = housekeeping gene. Graph represents the mean ± SD, **** p < 0.0001.

Both KO-BRCA1 pools exhibited elevated levels of total PR protein expression relative to the NE-1 control as well as evidence of multiple, basally phosphorylated protein species (upshifted bands; Figure 1A). Considering these observations, the R5020-mediated regulation of PR phosphorylation at multiple serine sites was investigated following a 1-hour R5020 treatment (Figure 1C). Interestingly, there was enhanced phosphorylation at phospho-PR serine sites S294 and S81 in the KO-BRCA1 pools, with the KO-B pool demonstrating the most pronounced increase. No differences in phosphorylation were observed at phospho-PR serine sites S345 and S190 between the KO-BRCA1 pools and the control. These findings suggest that BRCA1 depletion leads to basal and progestin-mediated hyperphosphorylation of PR-B at site-specific serine residues.

Additionally, to determine if BRCA1 depletion affected the transcriptional activity of PR, the expression of known PR-target genes following a 6-hour R5020 treatment were analyzed (Figure 1D). Both KO-BRCA1 pools, KO-A and KO-B, exhibited significantly higher mRNA levels of *PR* and *FOXO1* when compared to their respective vehicle control and the R5020-treated NE-1 control (p < 0.001). Notably, KO-B pool demonstrated the most robust transcriptional response, with significantly increased mRNA levels of *PR* and *FOXO1* when compared to the R5020-treated KO-A pool (p < 0.001). All samples exhibited elevated mRNA levels of *p21* mRNA expression after R5020-treatment when compared to their respective vehicle control (p < 0.001) but had marked differences between them. KO-A pool exhibited higher mRNA levels of *p21* when compared to the R5020-treated NE-1 control (p < 0.001), while KO-B exhibited decreased mRNA levels of *p21* when compared to the R5020-treated NE-1 control (p < 0.001). These findings suggest that BRCA1 depletion generally potentiates PR-mediated transcription, but certain targets such as p21 may be subject to other regulatory mechanisms.

### Migration and Cytoskeletal Organization are the predominant drivers of BRCA1-dependent/progestin-induced gene signature

To identify novel transcripts regulated by progestin signaling in the context of BRCA1 depletion, an RNA-seq study was conducted (Figure 2). The NE-1, and both KO-BRCA1 pools, KO-A and KO-B, were treated with equivalent volume vehicle (ethanol) or 10nM R5020 for 6-hours, RNA isolated and submitted for RNA quality analysis, library construction, and subsequent sequencing (University of Minnesota Genomics Center). Successful BRCA1 depletion was validated by observing a reduction of BRCA1 normalized transcript counts across both KO-BRCA1 pools, KO-A and KO-B (Figure 2A). Principle Component Analysis (PCA) demonstrated tight clustering of biological replicates, where the primary driver of variance was progestin treatment (85%), followed by BRCA1 status (7%) (Figure 2B). To isolate the impact of BRCA1 depletion on the progestin transcriptome, the R5020-regulated genes in the NE-1 control were compared against the combined R5020-regulated gene signature of both KO-BRCA1 pools, KO-A and KO-B (Figure 2C). The genes commonly regulated in the NE-1 control and across all lines were filtered out, and the non-overlapping KO-BRCA1 80 gene signature was analyzed. This non-overlapping cohort represents genes that are uniquely modulated by R5020 upon BRCA1 depletion, suggesting that BRCA1 interferes with progestin-dependent gene regulation when wild type levels of BRCA1 protein are present.

**Figure 2.**
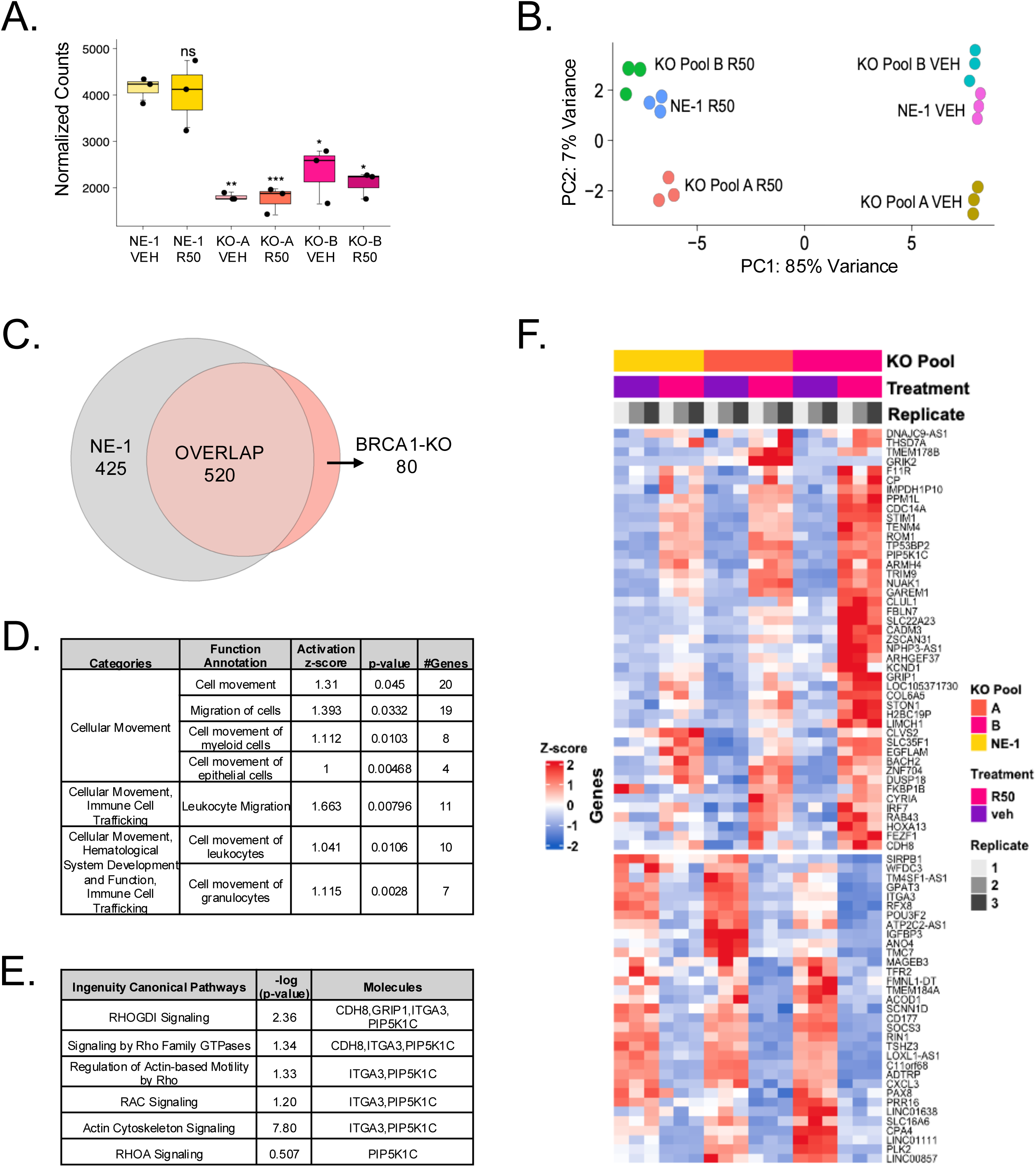
Migration and Cytoskeletal Organization are the predominant drivers of BRCA1-dependent/progestin-induced gene signature. **(A)** BRCA1 normalized transcript counts across the cell models. **(B)** Principal component analysis (PCA) for the cell models after treatment with vehicle (ethanol) or 10nM R5020 for 6 hours. Y-axis shows variance due to BRCA1 status; x-axis shows variance due to hormone treatment. **(C)** Euler diagram overlapping the R5020-regulated genes of NE-1 control vs. the combined R5020-regulated genes of both KO-BRCA1 pools, KO-A and KO-B. The BRCA1-dependent/progestin-induced gene signature is comprised of 80 genes. **(D)** Table of Ingenuity Pathway Analysis (IPA) identifying diseases and biological functions associated with the 80 gene signature. Table is comprised of categories, function annotation, activation z-score, p-values, and # genes associated with each function. **(E)** Table of IPA identifying canonical pathways associated with the 80 gene signature. Table is comprised of ingenuity canonical pathways, -log (p-value), and molecules associated with each pathway. **(F)** Heatmap of the 80-gene BRCA1-dependednt/progestin-induced signature. Data are clustered by row (gene) and column (replicate). Gene expression levels are represented as Z-scores, with red indicating high relative expression and blue indicating low relative expression.

To determine the biological impact of the BRCA1-dependent/progestin-induced 80 gene signature, transcriptomic data was analyzed using IPA for disease and biological function associations (Figure 2D) and established canonical pathways (Figure 2E). IPA revealed a significant enrichment of biological functions related to cellular motility, such as migration of cells, and cell movement of leukocytes, granulocytes, myeloid and epithelial cells (activation Z-score > 1). In addition, the most prominent canonical pathways identified in the 80-gene signature are those critical to structural remodeling, in particular, Rho Family GTPases signaling, Rac signaling, and actin cytoskeleton signaling. To visualize the expression dynamics of the BRCA1-dependent/progestin-induced 80-gene signature, a heatmap was generated comparing the NE-1 control to both KO-BRCA1 pools, KO-A and KO-B (Figure 2E). Gene expression values where normalized to Z-score to emphasize relative changes across replicates. The upper cluster contains genes upregulated by R5020, while the lower cluster contains genes that were downregulated. Within the 80-gene BRCA1-dependent/R5020-induced signature, several key mediators of cytoskeletal organization were identified such as PIP5K1C, ITGA3, and CDH8. These findings suggest that BRCA1 depletion results in an induction of cell motility related genes.

### BRCA1 dependent/progestin-induced gene signatured leads to worse OS and PFS in ovarian cancer dataset

To evaluate the clinical relevance of the BRCA1-dependent/progestin-induced 80-gene signature, patient survival outcomes were assessed using the Kaplan-Meier plotter ovarian cancer dataset (Figure 3) (39). High expression of the upregulated genes of the 80-gene signature predicts worse overall survival (HR = 1.26, p = 0.026; Figure 3A) and worse progression-free survival (HR = 1.55, p = 3.6E-6; Figure 3B) compared to those with low expression. These findings suggest that elevated expression of the genes within the 80-gene signature significantly correlates with unfavorable patient prognosis.

**Figure 3.**
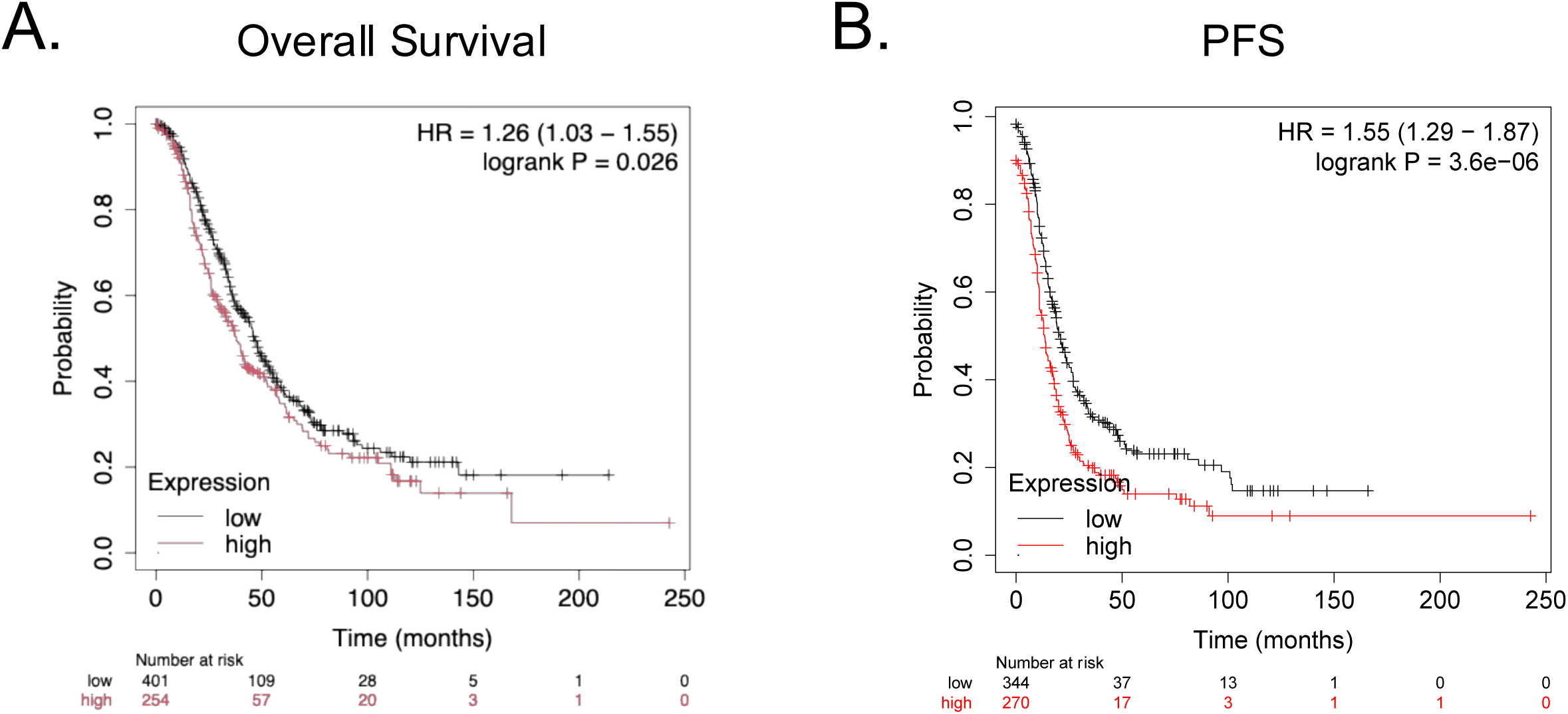
BRCA1 dependent/progestin-induced gene signature leads to worse OS and PFS in ovarian cancer dataset. **(A)** Kaplan-Meier (KM) for plot overall survival (OS) in the ovarian cancer dataset using the 80-gene BRCA1-dependent/progestin-induced gene signature. HR = 1.26 (1.03-1.55). logrank P = 0.026. **(B)** KM plot for progression-free survival (PFS) in the ovarian cancer dataset using the 80-gene BRCA1-dependent/progestin-induced gene signature. HR = 1.55 (1.29-1.87). logrank P = 3.6×10^-6^.

### BRCA1 depletion enhances progestin-driven migration

Since the RNA-seq analysis identified migration and cytoskeletal organization as the predominant pathways driven by the BRCA1-dependent/progestin-induced 80-gene signature, transwell migration assays were performed. Following a 24-hour R5020 pre-treatment and a subsequent 6-hour treatment at the time of transwell seeding, all cell lines exhibited significantly increased migration relative to their respective vehicle controls (NE-1, p = 0.004; KO-A and KO-B, p < 0.001) (Figure 4A-B). Notably, R5020-induced migration was significantly amplified in both KO-BRCA1 pools, KO-A and KO-B, when compared to the R5020-treated NE-1 control (p < 0.001). These findings suggest that BRCA1 depletion sensitizes cells to progestin-driven cell motility.

**Figure 4.**
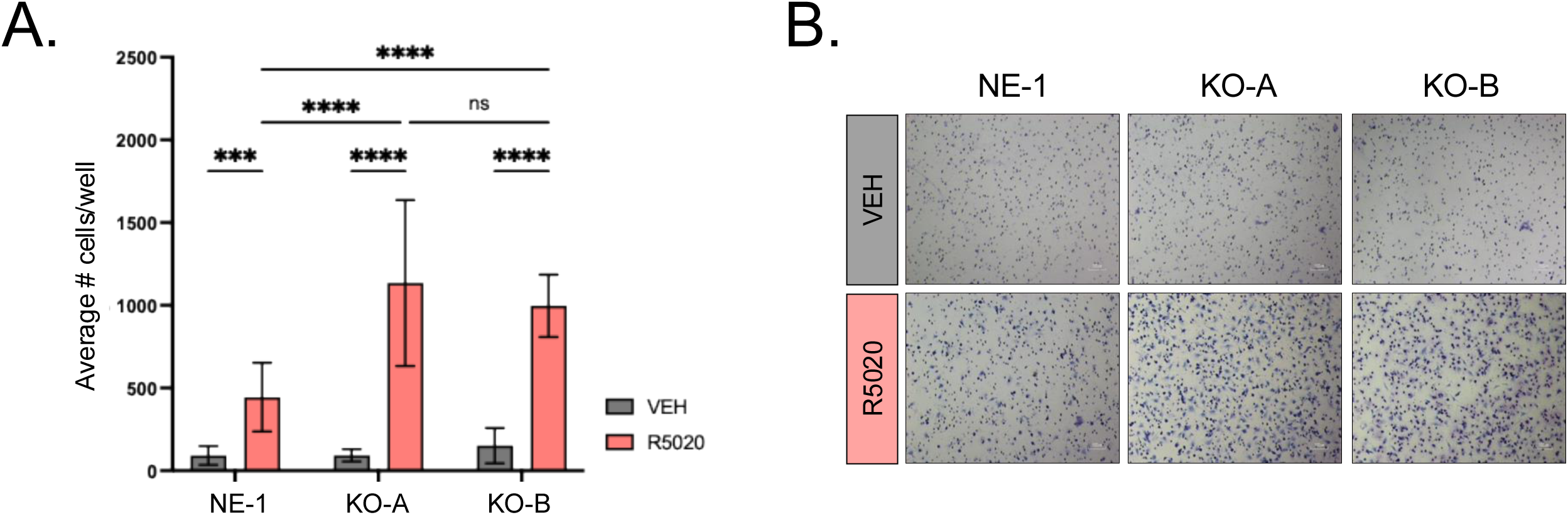
BRCA1 depletion enhances progestin-driven migration. **(A)** Average migration of NE-1 and KO-BRCA1 pools in transwell migration assay. Vehicle (ethanol) or 10nM R5020 treatment for 6-hour migration timepoint, which includes a prior 24-hour pre-treatment of either vehicle (ethanol) or 10nM R5020, respectively. 10% FBS used as chemoattractant. Graph represents the mean ± SD, *** p = 0.0004, **** p < 0.0001. **(B)** Representative bright-field images of A to the side. 100uµm magnification.

### Progestin-induced migration of PR+ ES2 cells is regulated by PIP5K1C/Rho/ROCK/Rac signaling

To investigate whether the potentiated R5020-induced migration observed in BRCA1-depleted cells was dependent on Rho-GTPase, Rac, and actin cytoskeleton signaling pathways identified by IPA (Figure 5A), transwell migration assays were performed using specific pathway inhibitors (Figure 5A-I). Following a 24-hour R5020 + PIP5K1C inhibitor (UNC3230) combo pre-treatment and a subsequent 6-hour treatment at the time of transwell seeding, UNC3230 significantly reduced R5020-induced migration in both KO-A (p < 0.0001) and KO-B (p = 0.0082) pools, while having no effect on the NE-1 control (Figure 5B). Notably, only in the KO-B pool did PIP5K1C inhibition restore migration to near-baseline levels, as KO-A remained significantly elevated (p = 0.0001) compared to its vehicle control. Following a 24-hour R5020 pre-treatment and a subsequent 6-hour R5020 + Rho Inhibitor (CT04) combo treatment at time of transwell seeding, CT04 significantly attenuated R5020-induced migration only in the KO-A pool (p = 0.0285) (Figure 5D). While a downward trend was observed in KO-B, it did not reach statistical significance. Notably, for both KO-BRCA1 pools, KO-A and KO-B, migration with the Rho inhibitor remained significantly higher to their own respective vehicle controls (p = 0.0095 and p=0.0092, respectively). These findings suggest that while PIP5K1C and Rho signaling contribute to the phenotype, they are not the primary driver of the potentiated cell motility seen upon BRCA1 depletion.

**Figure 5.**
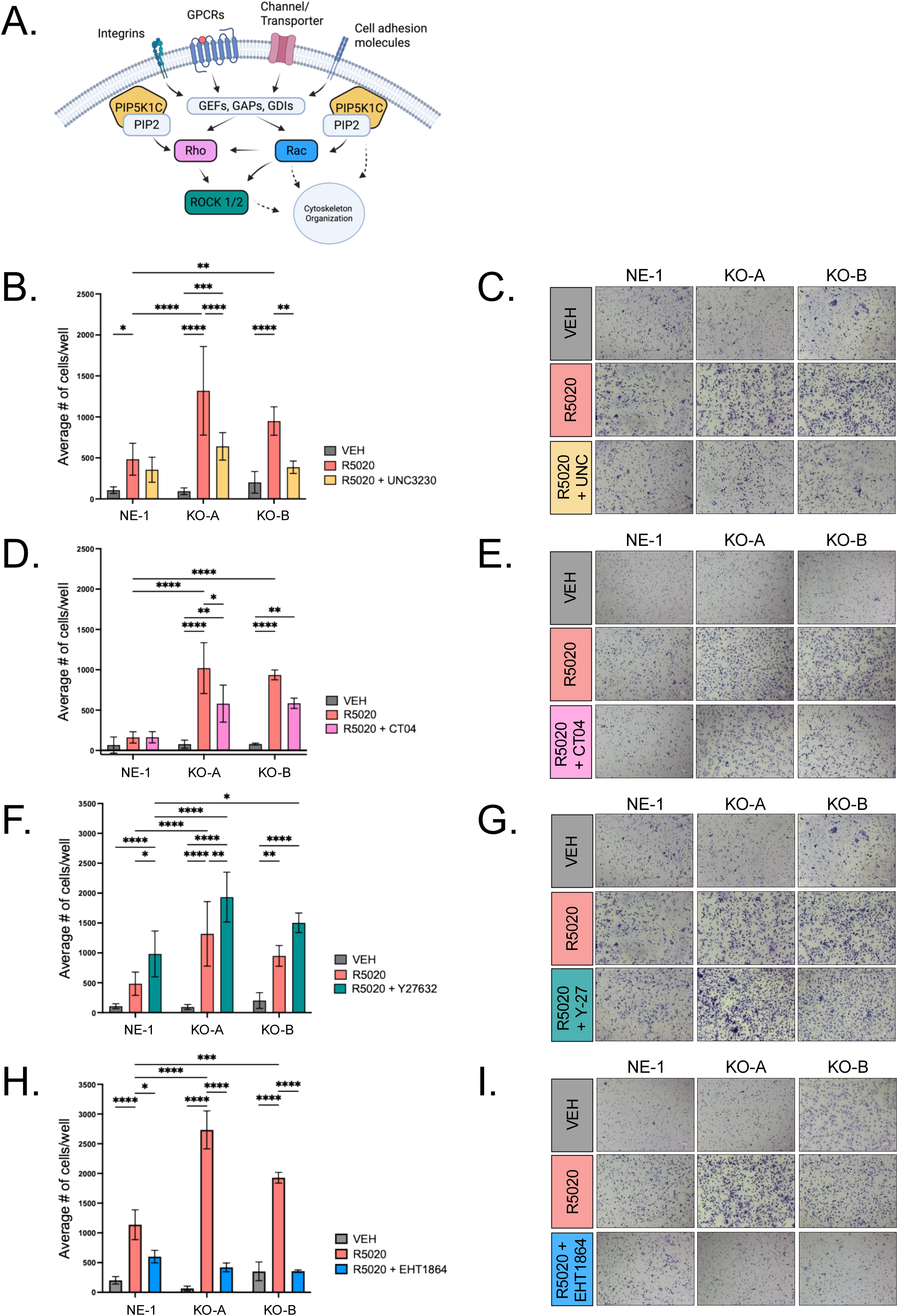
Progestin-induced migration of ES2 PR+ cells is regulated by PIP5K1C/Rho/ROCK/Rac signaling. **(A)** Signaling schematic representing the regulation by PIP5K1C/Rho/ROCK/Rac signaling. **(B-I)** Average migration of NE-1 and KO-BRCA1 pools in transwell migration assay. Vehicle (ethanol), 10nM R5020, or 10nM R5020 + inhibitor combo treatment for 6-hour migration timepoint, which includes a prior 24-hour pre-treatment of either vehicle (ethanol), 10nM R5020, or 10nM R5020 + inhibitor treatment, respectively. 10% FBS used as chemoattractant. Representative bright-field images to the side of each graph taken at 100µm magnification. **(B-C)** PIP5K1C Inhibitor: 10µM UNC3230. Graph represents the mean ± SD, * p = 0.0248, ** p ≤ 0.0082, *** p = 0.0001, **** p < 0.0001. **(D-E)** Rho Inhibitor: 1µg/ml CT04 (Note: no inhibitor pre-treatment). Graph represents the mean ± SD, * p = 0.0285, ** p ≤ 0.0095, **** p < 0.0001. **(F-G)** ROCK Inhibitor: 10µM Y-27632. Graph represents the mean ± SD, * p ≤ 0.0384, ** p ≤ 0.0016, **** p < 0.0001. **(H-1)** Rac Inhibitor: 25µM EHT1864. Graph represents the mean ± SD, * p = 0.0120, *** p = 0.0002, **** p < 0.0001.

Following a 24-hour R5020 + ROCK inhibitor (Y-27632) combo pre-treatment and a subsequent 6-hour treatment at the time of transwell seeding, Y-27632 significantly increased R5020-induced migration in NE-1 (p = 0.0198) and KO-BRCA1 pool, KO-A (p = 0.0014) (Figure 5F). A similar upward trend was observed in KO-B, but with no statistical significance. Furthermore, both KO-BRCA1 pools, KO-A and KO-B maintained significantly higher migratory capacity that the NE-1 control under ROCK inhibition (p < 0.001 and p = 0.0384, respectively). The fact that ROCK inhibition further increased rather than inhibited R5020-induced migration suggests that this specific signaling may operate under a negative feedback loop mechanism or that cells undergo a shift to a more migratory phenotype when ROCK is suppressed.

Following a 24-hour R5020 + Rac inhibitor (EHT1864) combo pre-treatment and a subsequent 6-hour treatment at the time of transwell seeding, EHT1864 significantly attenuated R5020-induced migration in NE-1 control (p = 0.0120) and both KO-BRCA1 pools, KO-A and KO-B (p < 0.0001) (Figure 5H). In all groups, the R5020 + EHT1864 inhibitor combination reduced migration to levels statistically indistinguishable from their respective vehicle controls. These findings suggest that the R5020-induced migration potentiated by BRCA1 depletion is primarily influenced by Rac signaling.

Overall, these findings suggest that BRCA1 depletion facilitates a pro-migratory state in an ovarian cancer cell model and is highly sensitive to progestin. The pro-migratory state is regulated by Rho-family GTPases, Rac, and actin cytoskeleton signaling pathways. PIP5K1C and Rho act as secondary modulators, ROCK signaling may work under a negative feedback loop mechanism, and Rac signaling serves as the essential driver of progestin-induced migration.

## DISCUSSION

BRCA1 is known to regulate progesterone receptor expression and transcriptional activity (15,25–27). Studies have reported that BRCA1 inhibits progesterone receptor (PR) activity by obstructing receptor recruitment to progesterone response elements (PREs) and promoting assembly of corepressor rather than coactivator complexes (40). Furthermore, BRCA1 is also shown to ubiquitinate PR using BRCA1/BARD complexes by targeting it for proteasomal degradation (27), a process that is lost in *BRCA1* mutation carriers. Consistent with the loss of this BRCA1 regulation, our results demonstrate that BRCA1 depletion leads to basal and progestin-mediated hyperphosphorylation of the PR-B isoform at site-specific serine residues, S294 and S81. This shift in the PR phosphorylation appears to functionally reprogram the receptor for enhanced genomic activity. This is evidenced by our observation that BRCA1 depletion potentiates PR-mediated transcription of key targets, including *PR* and *FOXO1*. Alternatively, while progestin-treatment increased *p21* mRNA, our results demonstrated that the response to BRCA1 depletion differed across both KO-BRCA1 cell pools. The observed variability suggests that *p21* may be potentially subject to other regulatory mechanisms than direct targets like *FOXO1*. This implies a degree of specificity in the PR-BRCA1 interaction.

Our transcriptomic analysis reveals that BRCA1 serves as a moderator of the progestin-regulated transcriptome. While progestin treatment was found to be the main driver of variance in the RNA-seq study, we were able to isolate an 80-progestin-induced gene signature uniquely modulated in the context of BRCA1-depletion, where migration and cytoskeletal organization where the predominant PR-regulated processes. This provides a link between progesterone receptor signaling and HGSOC cancer progression. Furthermore, the enrichment of biological functions related to cell movement and migration mirrors clinical observations in normal fallopian tube tissues from healthy *BRCA1/2* carriers, the cells of origin for HGSOC, which exhibit gene signatures similar to HGSOC only during the luteal phase of the menstrual cycle (28). This is supported by the activation of canonical pathways central to cytoskeletal reorganization, including Rho Family GTPases, Rac, and actin cytoskeleton signaling. Our identification of these pathways is corroborated by proteomic studies which demonstrated that BRCA1 deficiency in ovarian cancer is associated with the altered expression of key motility regulators, in particular those involved in Rho-GTPase signaling (41). By identifying specific genes within our 80-gene signature, such as PIP5K1C, ITGA3, and CDH8, we further define this pro-migratory phenotype.

By utilizing the Kaplan-Meier plotter ovarian cancer dataset, we established that high expression of the upregulated components of the BRCA1-dependent/R5020-induced 80-gene signature serves as a strong predictor of poor clinical prognosis.

Specifically, patients exhibiting high expression of this signature predicted significantly worse overall survival (OS) and progression-free survival (PFS). These clinical correlations confirm our in vitro findings, suggesting that the activation of the pathways central to cytoskeletal organization reflect a shift to an invasive and lethal disease state. Furthermore, the association of this signature with worse PFS suggests that this pro-migratory phenotype may facilitate treatment-resistance or promote rapid metastatic spread.

Progestins are known to regulate cell migration and invasion in breast cancer cell models through stabilization of the RhoA complex, regulation of key genes, and focal adhesion modulation (22,42,43). Consistent with this pro-migratory role, our results demonstrate that R5020 treatment significantly promotes ovarian cancer cell migration, and this migration is further enhanced in the context of BRCA1 depletion. This suggests that the 80-gene signature translates into a distinct invasive phenotype in BRCA1-deficient contexts.

The most significant finding of this study is the total attenuation of the R5020-induced migration by the Rac inhibitor EHT1864. BRCA1 is known to possess E3 ubiquitin ligase activity and can interact with various signaling proteins (27,44–46). Notably, interfering with the E3 ubiquitin ligase activity of BRCA1 protein leads to enhanced cell motility (46), connecting BRCA1-mediated ubiquitination to cell migration. Based on this information, it is possible that BRCA1 facilitates the degradation Rac-activating GEFs and, without BRCA1, progestin stimulation likely leads to an unchecked accumulation of active Rac-GTP. The observation that Rac inhibition was the only treatment to return all samples to baseline suggests that Rac is the primary effector driving R5020-induced cell migration in the context of BRCA1 depletion. Conversely, the partial migratory inhibition seen with PIP5K1C and Rho inhibitors suggests that these pathways are active but are not the major drivers for motility in this model. The finding that ROCK inhibition by Y-27632 further exacerbated R5020-induced migration is particularly striking. However, this is consistent with established signaling crosstalk, as inhibition of ROCK leads to Rac activation (47). In the context of BRCA1 depletion and progestin stimulation, inhibition of ROCK possibly acts though a negative feedback loop mechanism that leads to a Rac-dependent pro-migratory state.

Collectively, our findings indicate that the normal response to progesterone is disrupted in the absence of BRCA1. We show that BRCA1 depletion sensitizes ovarian cancer cells to progestin-induced migration by altering PR phosphorylation and remodeling cytoskeletal organization via PIP5K1C, Rho-GTPase, and Rac1-dependent mechanisms. Future studies utilizing targeted siRNA-mediated knockdown will be essential to substantiate these signaling mechanisms. By unraveling the PR-BRCA1-cell motility interaction, we may uncover novel opportunities for intervention and prevent the metastatic spread of HGSOC.

## ACKNOWLEDGEMENTS

The University of Minnesota Genome Engineering Shared Resource (GESR) provided the genome edited cell lines. The University of Minnesota Genomics Core (UMGC) performed the gene expression sequencing. This work was supported by the Tickle Family Land Grant Endowed Chair of Breast Cancer Research (to CAL) and a Translational Women’s Cancer Research Award (to CAL).

